# Low Dimensional Activity in Spiking Neuronal Networks

**DOI:** 10.1101/109900

**Authors:** Emil Wärnberg, Arvind Kumar

**Author notes:** Corresponding authors: Arvind Kumar.

## Abstract

Several recent studies have shown that neural activity *in vivo* tends to be constrained to a low-dimensional manifold. Such activity does not arise in simulated neural networks with homogeneous connectivity and it has been suggested that it is indicative of some other connectivity pattern in neuronal networks. Surprisingly, the structure of the intrinsic manifold of the network activity puts constraints on learning. For instance, animals find it difficult to perform tasks that may require a change in the intrinsic manifold. Here, we demonstrate that the *Neural Engineering Framework* (NEF) can be adapted to design a biologically plausible spiking neuronal network that exhibit low dimensional activity. Consistent with experimental observations, the resulting synaptic weight distribution is heavy-tailed (log-normal). In our model, a change in the intrinsic manifold of the network activity requires rewiring of the whole network, which may be either not possible or a very slow process. This observation provides an explanation of why learning is easier when it does not require the neural activity to leave its intrinsic manifold.

**Significance statement:** A network in the brain consists of thousands of neurons. A priori, we expect that the network will have as many degrees of freedom as its number of neurons. Surprisingly, experimental evidence suggests that local brain activity is confined to a space spanned by 10 variables. Here, we describe an approach to construct spiking neuronal networks that exhibit low-dimensional activity and address the question: how the intrinsic dimensionality of the network activity restricts the learning as suggested by recent experiments? Specifically, we show that tasks that requires animals to change the network activity outside the intrinsic space would entail large changes in the neuronal connectivity, and therefore, animals are either slow or not able to acquire such tasks.

## Introduction

The availability of novel experimental methods allows for simultaneous recording of tens to hundreds of neurons and has made it possible to observe the fine structure of temporal evolution of task-related neuronal activity *in vivo*. The multi-unit neuronal activity can be described in terms of an *N* dimensional neural state-space where each axis (typically) corresponds to the firing rate of each neuron. The activity at a particular time corresponds to a point in this space, and the temporal evolution of the neuronal activity constitutes a trajectory. Analysis of such trajectories has revealed that across different brain regions and in different behavioral conditions the neural activity remains low dimensional (Mazor and Laurent, 2005; Ganguli et al., 2008; Cunningham and Yu, 2014; Sadtler et al., 2014; Mazzucato et al., 2016; Williamson et al., 2016; Murray et al., 2016). That is, the trajectories corresponding to the task-related activity tend to be constrained to a hyperplane (“intrinsic manifold”) in the state space rather than moving freely in all directions.

Treating the brain as a dynamical system, the structure of the activity trajectories in the neural space determines the function of the neuronal network. This was best illustrated by an experiment involving brain-computer-interface (BCI) learning in monkeys. Sadtler et al. (2014) showed that animals were able to quickly learn the BCI task when the neural activity mapping was confined to the intrinsic manifold of the activity. By contrast, BCI learning was slow when the neural activity mapping for the BCI task was outside of the intrinsic manifold Sadtler et al. (2014). These observations raise two pertinent questions: (1) what is the origin of low-dimensional activity in spiking neuronal networks and (2) why is it difficult to learn outside manifold mapping or to alter the structure of the intrinsic dynamics?

A homogeneous balanced random recurrent network which have been very successful in modeling the statistics of spiking activity and pairwise correlations *in vivo* (Brunel, 2000; Kumar et al., 2008) cannot generate low-dimensional activity (Mazzucato et al., 2016; Williamson et al., 2016) because the balance of excitation and inhibition actively decorrelates the population activity (Renart et al., 2010; Tetzlaff et al., 2012). Therefore, clustered architecture has been proposed, in which the dimensionality of activity is defined by the cluster count (Mazzucato et al., 2016; Williamson et al., 2016). However, there is no direct evidence of clustered architecture in the neocortex (Boucsein et al., 2011; Schnepel et al., 2015; Perin et al., 2011; Jiang et al., 2015). In fact, even at the functional level neurons cannot be separated among distinct clusters (Williamson et al., 2016). Finally, even though the cluster-based network architecture explains how the dimensionality of the activity may depend on the size of the neuron population, it does not explain why it should be difficult to change the dynamics outside the intrinsic manifold. Thus, it remains an open question to identify underlying mechanisms that can generate low-dimensional activity.

Here, we show that it is possible to design non-clustered spiking neuronal networks that can generate low-dimensional activity. We found in such a network neurons receive balanced excitation and inhibition, and synaptic weights show a heavy-tailed distribution. This model suggests that small changes in the weight matrix are sufficient to generate a new activity that lies within the intrinsic manifold, whereas nearly all the synapses have to be altered to generate activity that lies outsides the intrinsic manifold. Learning such large-scale changes in the weight matrix are likely to be slow, and, therefore, animals may find it difficult to learn tasks that involve generation of activity outside the intrinsic manifold.

## Methods

### Dynamical system specification

The core of our approach to design spiking neuronal networks whose activity is confined specific intrinsic manifold involves mapping the spiking activity to a prescribed dynamical system. To this end we used the *Neural Engineering Framework* (NEF) (Eliasmith and Anderson, 2003) to design a spiking neuronal network with approximately four dimensional dynamics. The approach is general and can be used to design a network with any arbitrary dimensional dynamical system. The NEF, like many of the similar models (e.g. Boerlin et al., 2013), requires a dynamical system to describe the evolution of the fictive currents (see Eq. 7), or encoded values. One popular choice is to use a straight-forward integrator (see e.g. Eliasmith, 2005; Boerlin et al., 2013; Hoerzer et al., 2012). In the absence of input, such a network will retain whatever spiking rates it is set at. Such a network would contradict experimental evidence (Shafi et al., 2007), and stationary firing rates would by definition have no variance and hence be zero-dimensional. Therefore, we designed our dynamics following the perhaps secondly most popular choice, namely, an oscillating system (similar dynamics have also been used by e.g. Eliasmith, 2005; Denève and Machens, 2016).

Furthermore, in order to demonstrate the possibility of creating a network with low dimensionality as clearly as possible, we wanted the fictive currents (see Eq. 7) each to have approximately equal variance. Had the variance of the underlying dynamical system been unequally distributed among its variables, the four dimensions found by the PCA would most likely also have had unequal variance. For the same reason, we wanted all the fictive currents to be uncorrelated. To comply with these requirements, we designed the dynamics so that the first two encoded values would be the leading and lagging components of a 2 Hz oscillator and the last two would similarly be the lead and lag of a 4 Hz oscillator. Additionally, we added a non-linear term causing the amplitude to converge to the normalized value 1. With these terms combined, the differential equations for the dynamical system become:

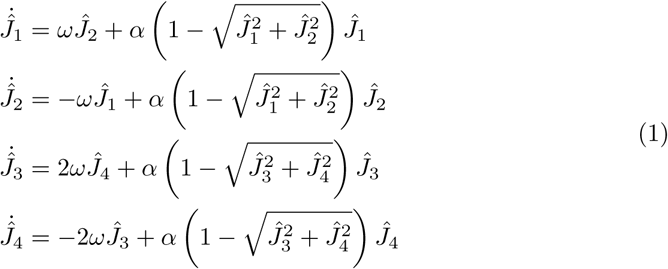

We used *α* = 0.2 which is enough to help stabilize the dynamics. An advantage of having the regularization term is also that the network neither requires a particular starting state nor any particular external input. This helps support the claim that the network is truly autonomously generating low dimensional activity. However, this can be altered and it is possible to adapt the network such that a specific input drives the network to the specific manifold.

We chose to encode an oscillator as the underlying dynamical system for its computational simplicity and any other dynamics can also be used without affecting the main results. Indeed, having the fictive currents endlessly stuck in a simple limit cycle is rather uninteresting from the perspective of using them to encode behaviorally relevant variables. However, Churchland et al. (2012) investigated the dynamics in the activity of the primate motor cortex, and found a strong oscillatory component when the activity was projected down on the first few principal components. Therefore, there is some tentative support for believing our choice of fictive currents dynamics to be reasonably representative of actual dynamics. Furthermore, note that *oscillator* here refers to oscillations of the encoded values, not of the neural activity directly.

### Estimation of network connectivity

To create the network structure, we used the NEF as implemented in the simulator package Nengo (Bekolay et al., 2014) (version 2.2.1). Following the specification of a dynamical system, Nengo creates a set of representative samples of the spike trains. In Nengo these samples are called *evaluation points,* and each sample corresponds to one instance of a set filtered spike trains *u_j_* (*t*) at a particular point in time. In our network, the number of such samples was 10,000. For each of these samples, the corresponding set of desired currents can be derived from the provided dynamical system. In other words, this is a set of relationships of the type *when the filtered incoming spike trains are like this, the input currents should be like that, if the encoded values are to evolve as prescribed.* To fit the weights, in its default state, Nengo performs an ordinary linear least squares optimization. However, as described in the **Results**, we modified this procedure so that biological constraints could be imposed on *W*. Technically, we implemented our modification as a subclass of the `Solver` class in Nengo. Our subclass takes as input a *connectivity matrix* where the elements are 1, −1 or 0, representing excitatory, inhibitory and absent synapses respectively. When called from Nengo, it used the `scipy.optimize` routine `nnls` to find a weight matrix minimizing the *L*2 decoding error while conforming with the connectivity matrix. Note that for creating Fig. 7B, the same connectivity matrix was used across all the simulations. Additionally, for Fig. 7B, the number of evaluation points was increased to 40,000 to decrease the variance.

### Simulation parameters

Neurons were modeled as Leaky-Integrate-and-Fire (LIF) units with somatic time constant 20 ms and an absolute refractory time of 2 ms. The post-synaptic currents were modeled as exponentially decaying with time constant 10 ms for both excitatory and inhibitory synapses.

The NEF and its implementation in Nengo require the specification of a few more parameters. One such parameter is the encoder matrix *K* (see Eq. 6). Following the default setting in Nengo, we chose this matrix randomly, such that the preferred direction of each neuron, as specified by the corresponding row of *K*, was drawn from a uniform distribution. Each row was however scaled (the scaling factor is called *gain* in Nengo) so that the maximal firing rates of the neurons are between 80 Hz and 120 Hz. Each neuron was also given an independent static bias current. These were randomly drawn from a distribution defined so that the so called *intercepts* become uniformly distributed. Intercepts can be understood as the smallest synaptic currents required for the neuron to spike. The bias currents cause the network to be spontaneously active, which removes the need for an external drive.

### Dimensionality estimation

The literature on low dimensional neural activity is mostly split between using PCA (e.g. Hoerzer et al., 2012; Churchland et al., 2010; Mazzucato et al., 2016; Murray et al., 2016) and *Factor Analysis* (FA, e.g. Sadtler et al., 2014; Williamson et al., 2016). As described by Cunningham and Yu (2014), there are only a few differences between the two methods. Most prominently, FA assumes that the variance of each neuron consists of one individual component and one shared. Hence, fluctuations originating from internal noise can be separated from the actual population signal. However, one crucial restriction with FA is that the dimensionality needs to be assumed a priori. Although an estimation can be obtained for example by cross-validation, it anyhow impedes the possibility of showing an unbiased distribution of the variance across the principal components. For this reason, we have only used PCA in this work.

## Results

There are two natural candidate mechanisms that may lead to a low-dimensional activity in a recurrent network. First, the network is driven by a low-dimensional external input. Second, the activity may reflect the intrinsic structure of the behavioral tasks that the network has learned, that is, the recurrent inputs are themselves low-dimensional. For instance, given the muscle synergies and laws of motion, most motor actions are intrinsically low-dimensional (Ingram et al., 2008; Thakur et al., 2008). Here, we explore the second possibility and provide an explanation of why it may be difficult to learn outside-manifold mappings in a BCI task. To this end we first provide a systematic approach to construct a spiking neuronal network with a prescribed dimensionality and dynamical system.

### Neuron transfer function does not increase the dimensionality of the output

When discussing dimensionality of neural activity, there are several physical quantities that could be assigned to the abstract concept of “activity”. Three reasonable choices are (a) firing rate, (b) sub-threshold membrane voltage and (c) total synaptic activity or, equivalently, total input current to a neuron. Given that neurons can be non-linear systems, it is necessary to establish how the dimensionality of one of these measures of neural activity translates to the dimensionality of others. Specifically, if we can establish that the dimensionality of the input current (or sub-threshold membrane potential) leads to the same or lower dimensionality in the spiking activity, it would make it easier to design a network with a low-dimensional activity.

To develop an intuition about this issue, we stimulated an ensemble of 1000 LIF neurons with a two dimensional input current and measured the response in spiking rates. We omitted any recurrent connections among the neurons in order to isolate the effect of the neuron transfer function from any transformation resulting from the network structure (Fig. 1A). The neurons were realized as firing rate units or with the spiking leaky-integrate-and-fire (LIF) model. For the firing rate model, to map the input current to output firing rate we considered four different transfer functions (linear, threshold linear, logistic function and step function, see Fig. 1B, top). All neurons in the ensemble had identical transfer functions. The LIF-neuron explicitly generates spikes upon reaching a spike threshold. Therefore, to estimate the firing rate, we counted spikes from each neuron in 40 ms time bins. To make the rate models comparable we averaged the rate models over 40 ms intervals.

**Fig 1.**
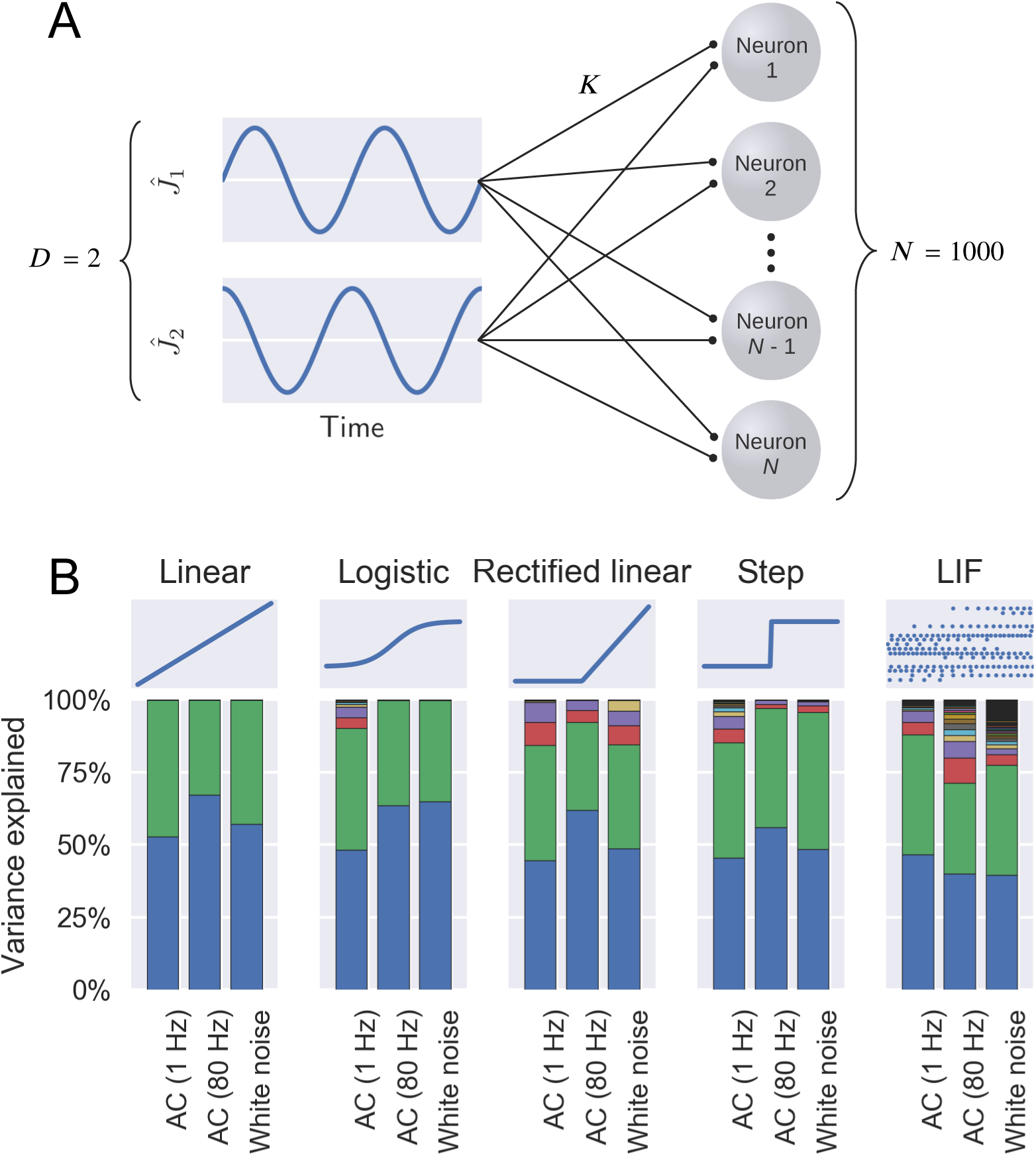
The resulting dimensionality after applying somatic transfer functions. (**A**) Schematic representation showing the sampling of a 2-dimensional input by an ensemble of unconnected neurons (see Eq. 2). (**B**) The output firing rates were analyzed using PCA. Each bar describes the distribution of the variance along the principal components for the respective combination of firing rate model (show on top) and input current shape (shown at the bottom). Each color in a bar shows the proportion of variance explained by the respective principal component. As is evident in every case only two principal components are sufficient to explain most of the variance.

The neuron ensemble received a two-dimensional input composed of two orthogonal time varying signals *ĵ*_1_(*t*) and *ĵ*_2_(*t*), and each neuron sampled the two components with synaptic weights 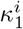 and 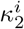. That is, each neuron *i* received an external current

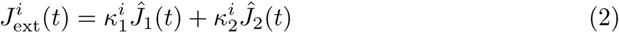

The scalar coefficients 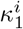 and 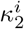 were drawn from a normal distribution independently for each neuron. We considered two different temporal dynamics for *ĵ*_1_(*t*) and *ĵ*_2_. First, we let *ĵ*_1_(*t*) = cos(2*πft*) and *ĵ*_2_(*t*) = sin(2*πft*) with *f* = 1 Hz or 80 Hz. Second, we let both *ĵ*_1_(*t*) and *ĵ*_2_(*t*) to be Gaussian white noise and their respective value at each time was drawn from two independent normal distributions with zero mean.

We used Principal Component Analysis (PCA) to determine the effective dimensionality of the ensemble output firing rate for each input. The PCA suggested that there are clearly two prominent dimensions for all combinations of **ĵ**(**t**) and neuron models tested (Fig. 1B). Although the non-linearities in the neuron transfer-function tended to distort the dimensionality of the neuron ensemble as expected, the contribution of the additional dimensions remained very small. Notably, even when we explicitly modeled the spiking behavior the dimensionality of the input currents was preserved both in the mean driven (sinusoidal input) and the fluctuation driven (white noise input) regime. These results also show that the spectrum of the input does not play a prominent role in the transfer of dimensionality as both narrow-band sinusoidal inputs and broad band white noise resulted in similar dimensionality in the output.

These results clearly suggest that a spiking neuronal network, in which the total (i.e. external and synaptic) input to neurons is low dimensional, will also have a low dimensional spiking or firing rate pattern, for different types of biologically plausible neuron transfer functions. The task of synthesizing a network with low dimensional *activity* can therefore be reduced to the problem of how to constrain the *input currents* to a low dimensional manifold.

## Generation of low dimensional input currents

The synaptic input current to neuron *i* can be written as

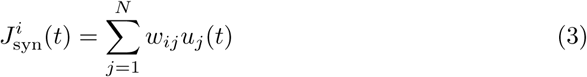

where *w_i,j_* is the synaptic weight between the post-synaptic neuron *i* and a pre-synaptic neuron *j*, and *u_j_*(*t*) is the filtered version of the spike train originating from neuron *j*:

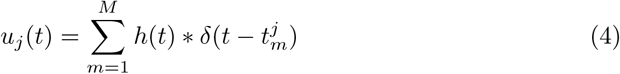

where *h*(*t*) is the synaptic kernel and 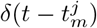 represents a spike at time *t_m_* from neuron *j*. For the immediate discussion, we remain agnostic about the exact shape of synaptic kernel *h*(*t*) (in our simulations described below, we used exponential kernels with time constant 10 ms). In matrix notation, Eq. 3 can be written as

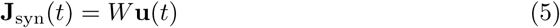

If *W* has rank *r*, **J**_syn_(*t*) will be constrained to a subspace with dimensionality at most *r* — in particular, the subspace spanned by the columns of *W*. If *W* is heavily degenerate so that *r* ≪ *N*, the input currents will be low dimensional. However, *W* having low rank does not in itself enable control over the shape of the input currents. Additionally, having a degenerate weight matrix is a synthetic global property which may be hard to enforce locally by individual synapses. In the following sections, we describe how to create and control low dimensional currents, first with *W* having low rank but then also how to relax that requirement.

### Controlling the dynamics of synaptic inputs in a recurrent network

Assuming **J**_syn_(*t*) has dimension *D*, its components can by definition be written as

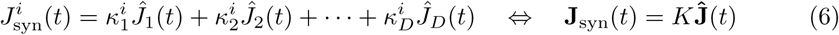

for some *fictive currents ĵ*_1_(*t*),…, *ĵ*_D_(*t*). For *D* = 2, Eq. 6 is similar to Eq. 2, with the important distinction that Eq. 2 describes an externally imposed current while the current in Eq. 6 is the current originating from the synaptic connectivity within the network. The question now is whether this intrinsic **ĵ** can be designed to evolve according to some specified dynamics?

Because the relationship between **J**_syn_(*t*) and **ĵ**(*t*) is linear, **ĵ** (*t*) can also be written as a linear combination of the filtered spike trains,

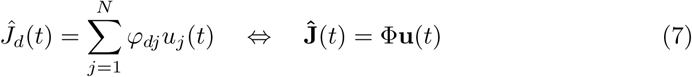

for some other coefficients {*φ_dj_*}. We can use the *Neural Engineering Framework* (NEF, Eliasmith and Anderson, 2003) and choose these coefficients by applying a straight-forward linear least squares optimization, fitting the filtered spike trains onto a desired fictive current. This method becomes even more useful when we recall that the filtered spike trains *u_j_*(*t*) in turn are generated by the synaptic currents, which implies that **ĵ** can be chosen to be a function of the current firing rates of the neurons, and hence, implicitly, itself (see Fig. 2). This implies that **ĵ** can be specified for a set of differential equations and not just a fixed trajectory.

**Fig 2.**
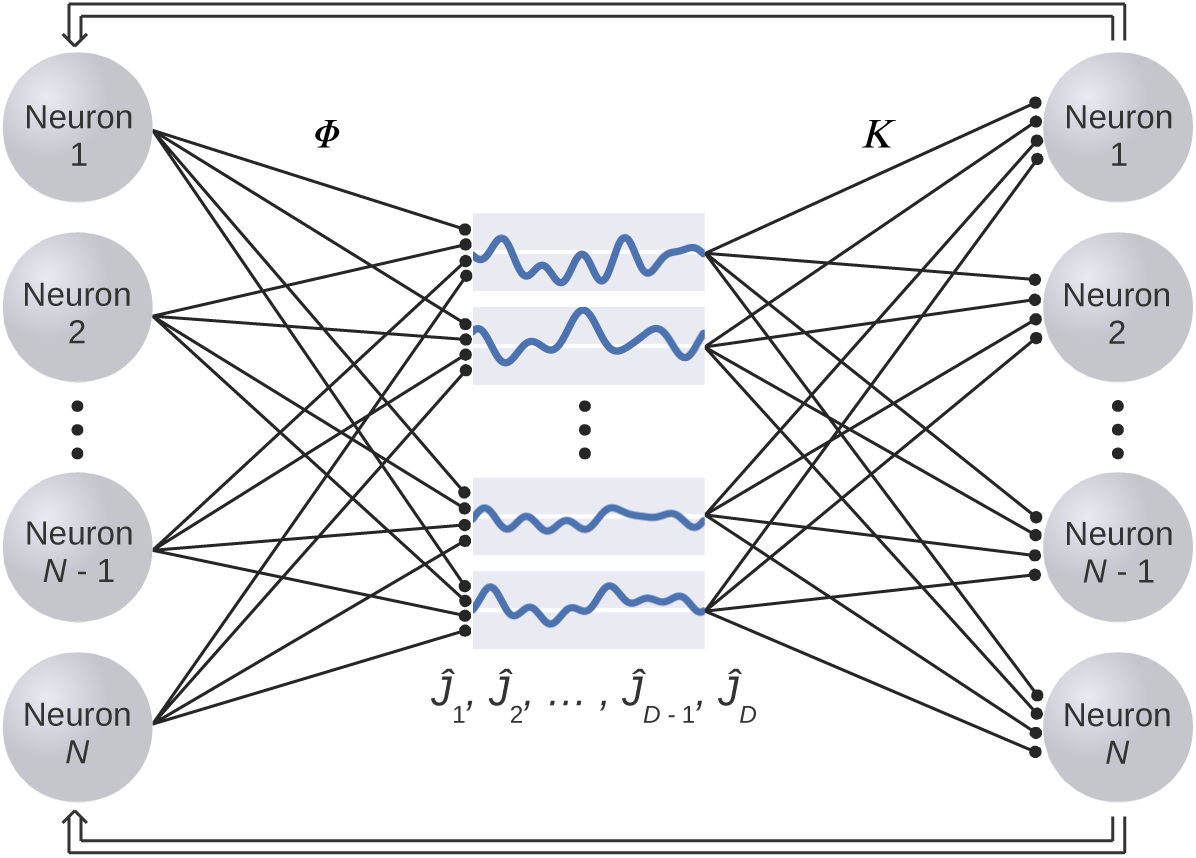
Schematic description of the procedure to design networks with a prescribed dimensionality. Here we have factored the weight matrix into two components. The weight matrix Φ transforms the filtered spike trains to the fictive currents **ĵ** (Eq. 7). The fictive currents constitute the low-dimensional input that in turns drive *the same* neuron population, with a scaling *K*. Note that the separation of the network in three layers is only to describe the procedure and there is only one set of neurons. The eventual recurrent connectivity is then described as *W* = *K* Φ.

Combining Eq. 6 and 7, we identify the similarity with Eq. 5:

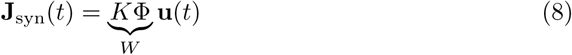

Because *K* ∈ ℝ^*N*×*D*^, Φ ∈ ℝ^*D*×*N*^ and *W* = *K*Φ, the rank of *W* will be at most *D*. As mentioned above, this is not obviously desirable. For this reason, Fig. 2 should be seen as strictly conceptual and not directly corresponding to any anatomical structure — the neurons are actually connected using a weight matrix *W* = *K*Φ, as in Eq. 3. In particular, *K* and Φ are only used to choose a suitable *W*.

## Biologically constrained weight matrix

While the matrix *W* ensures that the network will have low-dimensional activity, the above derivation of the weight matrix does not impose any constraints on *W* through *K* and Φ. In particular, biological weight matrices are typically sparse and adhere to *Dale’s law* (Strata and Harvey, 1999), i.e., that all the outgoing synapses from a neuron are either excitatory or inhibitory. A few elegant tricks have been proposed for separating the population into excitatory and inhibitory fractions (Parisien et al., 2008; Boerlin et al., 2013). Here we chose a more straight forward approach: rather than finding the optimal Φ, we applied the least square optimization to each neuron individually. This meant the coefficients found by the solver directly corresponded to the elements in *W*, and constraints imposed on them transferred directly to the weight matrix.

Using the above mentioned approach we constructed a network with *N* = 5000 neurons and *D* = 4 fictive currents, and constrained the linear least squares solver to find a weight matrix where the post-synaptic weights of the first 4000 (i.e., 80%) neurons are non-negative, and the last 1000 (i.e. 20%) are non-positive. This approximated the fraction of excitatory and inhibitory neurons found in the primate cortex (Braitenberg and Schüz, 1998). Additionally, to impose the constraint of sparse connectivity, we randomly chose 75% of the elements of the weight matrix to constrain to zero. The weight matrix resulting from this procedure can be seen in Fig. 3A. Note that a synapse might well be zero without being constrained to zero. Intuitively, this can happen if the least-squares optimal weight for an excitatory synapse would have been negative, or vice versa. Biologically, one might relate them to *silent synapses.* Because of this effect, the probability of an active connection was 10.77% between excitatory neurons, 10.85% from excitatory to inhibitory, 18.89% from inhibitory to excitatory, and 18.76% between inhibitory neurons. These connection probabilities are well within the biological ranges (Boucsein et al., 2011; Kätzel et al., 2010; Petreanu et al., 2009). Furthermore, because the set of possible presynaptic neurons is chosen independently for each postsynaptic neuron, the weight matrix can no longer be decomposed into *K* and Φ, and its rank is not strictly bound by *D*. Indeed, the singular values of the weight matrix shown in Fig. 3B indicate that its rank is larger than *D* = 4 for our example problem.

**Fig 3.**
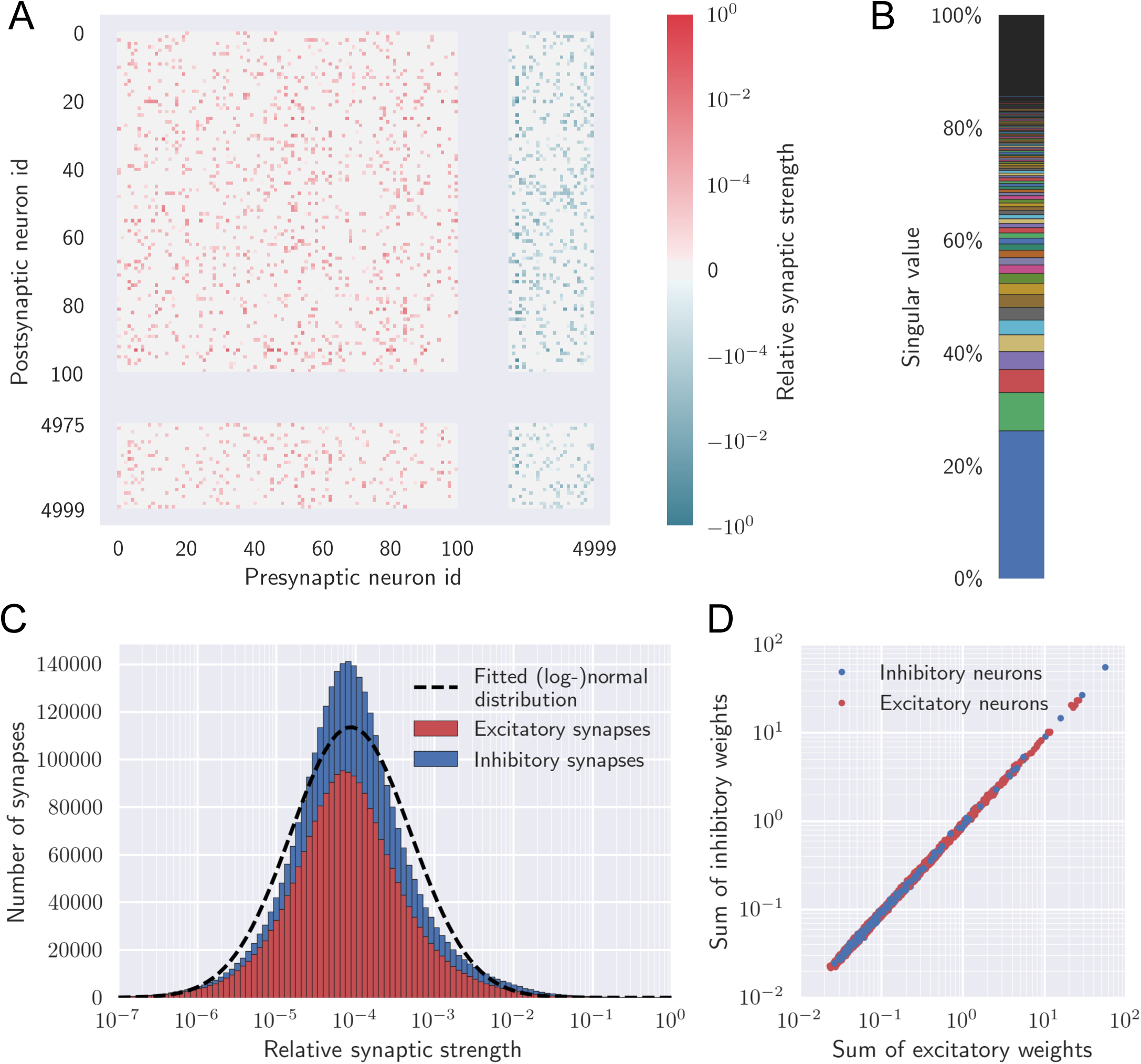
The weight matrix. (**A**) The color indicates synaptic weight, the scale is logarithmic on both sides of zero respectively. Red indicates excitatory and blue indicates inhibitory synapses. (**B**) The singular values of the weight matrix. The values are tilted, but the matrix clearly has higher rank than 4. (**C**) The distribution of values in the weight matrix. Note that the distribution definitely is heavy-tailed albeit not perfectly log-normal. (**D**) Summation of all incoming excitatory and inhibitory synaptic weights to each neuron (one row of *W*). The total weight of excitatory synapses (abscissa) matches the total weight of inhibitory synapses (ordinate) for each neuron.

The Daleian and the sparse connectivity were both forced into the design of the weight matrix. However, an interesting property emerged without the addition of any explicit constraints, namely, a heavy-tail on the distribution of synaptic weights (Fig. 3C). Although a closer analysis reveals it to be somewhat leptokurtic (kurtosis is 4.56 for logarithmized weights), the weight distribution is in any case rather close to being log-normal across more than five orders of magnitude. This conforms with recent experimental data that suggests that the weight distributions in biological neural networks are typically heavy-tailed, if not perfectly log-normal (Buzsáki and Mizuseki, 2014). In addition, the sum of excitatory and inhibitory weights is ≈ 0 for each neuron (Fig. 3D). In summary, even though we have used an engineering based approach, the resulting weight matrix *W* conforms with several key properties of the synaptic connectivity *in vivo*.

## Low-dimensional activity in a spiking neuronal network

To confirm that indeed the network we have designed has a low-dimensional intrinsic activity dynamics we simulated a spiking neuronal network with 5000 LIF neurons connected according to the weight matrix shown in Fig. 3A. In order to reduce spike train regularity, independent white noise was added to each neuron. As can be seen in Fig. 4C and 4D, the firing pattern is neither perfectly regular nor perfectly irregular (Poisson), but in between.

**Fig 4.**
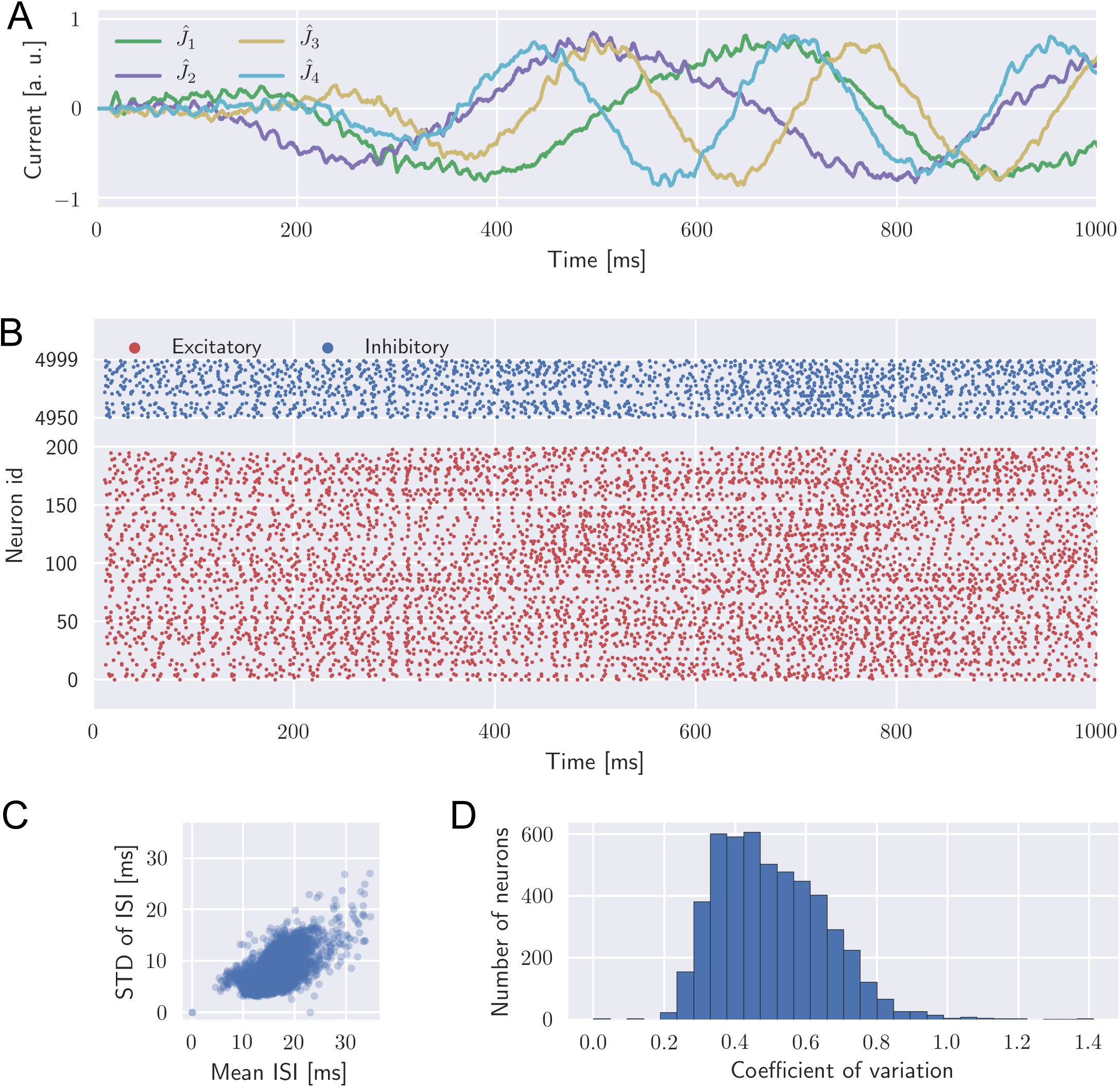
Low-dimensional network activity. (**A**) The four fictive currents (encoded values) during the first second of the simulation. (**B**) Spikes emitted by a selection of 250 neurons (200 excitatory (red) and 50 inhibitory(blue)) during the first second of the simulation. (**C**) The mean inter-spike interval (ISI) versus the standard deviation of the inter-spike interval for each neuron. Most of the points lie below the unity slope line indicating that non-Poissonian nature of the spiking activity. Inter-spike intervals longer than 100 ms were rare and have been excluded from the analysis. (**D**) Distribution of coefficients of variance of ISI.

The fictive currents we used to construct the network connectivity do not directly correspond to any measurable quantity in the network. Therefore, in order to verify that the network followed the instilled dynamic, we created four independent readout units. For the readout units, we found a set of weights Φ_readout_ using a linear least-squares optimization, similar to how the weights *W* were set. However, because these readouts do not correspond to any biological entity, we did not enforce the biological constraints for Φ_readout_. The readout values for the fictive currents can be seen in Fig. 4A. Note that they vary rhythmically. This is because the dynamical system we used to describe the fictive currents consisted of two limit cycles, with *ĵ*_1_ and *ĵ*_2_ oscillating at 2 Hz, and *ĵ*_3_ and *ĵ*_4_ at 4 Hz (see Methods).

The representational precision of the network is quite low, which can be seen in the wobbling of the fictive currents in Fig. 4A. In particular, for a network with 5000 neurons, it is considerably less precise than would be expected of similar models previously proposed (Eliasmith, 2005; Denève and Machens, 2016; Abbott et al., 2016). This is because of a combination of the added noise and the constraints in the solver. Specifically, the sparse connectivity implies that each neuron can only use the input from a little more than 10% of the network to estimate and calibrate its input current. Additionally, we used a synaptic time constant of only 10 ms, meaning the synaptic currents were not heavily smoothed and therefore were more difficult to linearly combine into smoothly varying fictive currents. Larger networks with slower synaptic time constants (e.g. those based on NMDAR and GABAB) would result in more precise fictive currents.

The purpose of our model is in any case not to improve coding or representational accuracy, but to demonstrate that it is possible to control the dimensionality of the network. To verify that the activity is indeed four-dimensional, we employed the same procedure we used to create Fig. 1. Namely, we divided the 10 s simulation duration into 250 consecutive bins, counted the number of spikes from each neuron in each bin and finally applied PCA on the sequence of spike count vectors (Fig. 5). The first four dimensions constitute 80% of the total variance (Fig. 5B), leading us to conclude that our network is essentially four-dimensional. These four dimensions are however not perfectly matched to the four fictive currents one-to-one (compare Fig. 4A and Fig. 5A). In other words, PCA does not fully discern the four “underlying” signals. Rather, the network activity projected down to the four first principal components evolve as a linear combination of the the fictive currents. For instance, even though we designed our network to exhibit two different limit cycles, the limit cycles constructed from the network activity are not perfect circles (Fig. 5C,D) but the periodicities at 2 Hz and 4 Hz are nonetheless clearly visible (Fig. 5A).

**Fig 5.**
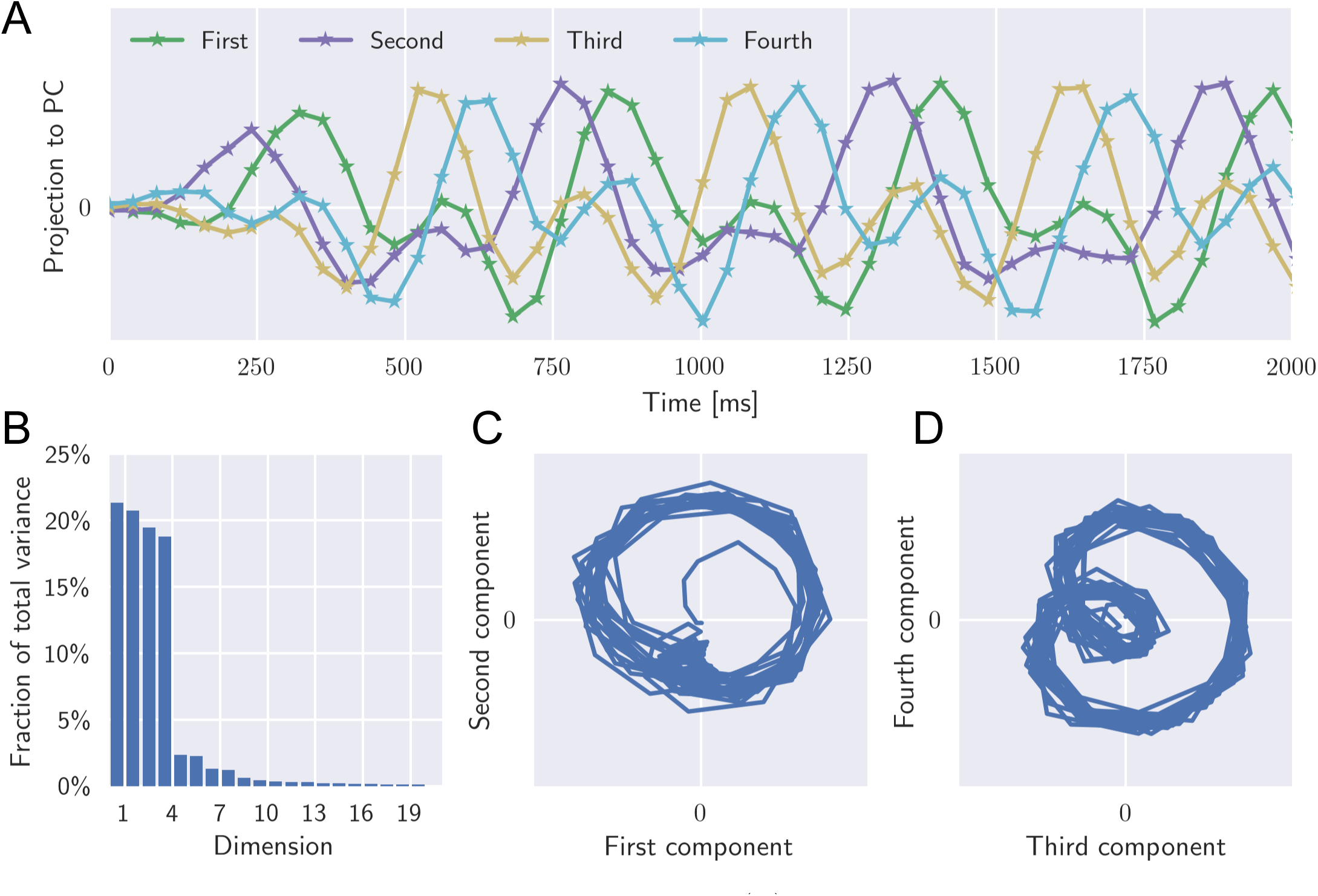
Principal component analysis of the spiking activity. (**A**) The first four principal components of the network spiking activity. (**B**) The fraction of the total variance explained by each principal component. Note that the activity is essentially four-dimensional, as intended. (**C**) and (**D**) Phase plots of the network activity (the same data as in A) indicating the underlying dynamical system that we intended to implemented in the network.

## Robustness of the low-dimensional activity manifold

The approach described above suggest that achieving a low dimensional activity dynamics requires fine tuning of the synaptic weights with a very high precision that may be biologically implausible. Moreover, synaptic weights are not likely to stay fixed in the brain *in vivo.* Stochastic nature of the vesicle release, activity dependent plasticity, stochastic changes in the spines, neuronal excitability and neuromodulators can perturb the synaptic strength at various time scales. MacNeil and Eliasmith (2011) found that perturbation of the synaptic weights can impair the stability of the attractor dynamics a network is designed to follow, but argued that synaptic learning rules can be designed to counter the effect. In any case, for low dimensionality, maintaining within-attractor dynamics is less important than maintaining the attractor itself. In other words, the temporal dynamics of the fictive currents in Fig. 2 are less important than maintaining the overall architecture.

To verify the robustness of the low dimensional activity to perturbations in synaptic weights, we scaled each synaptic weight by a factor *c_ij_* which was drawn from a normal distribution with mean 1.0 and standard deviation *σ* (Fig. 6A). Note that weight perturbations are not equivalent to addition of noise to each neuron. The stability of the low-dimensional activity was estimated in terms of the explained variance ratio of dimensions of the activity (analogous to Fig. 5B).

**Fig 6.**
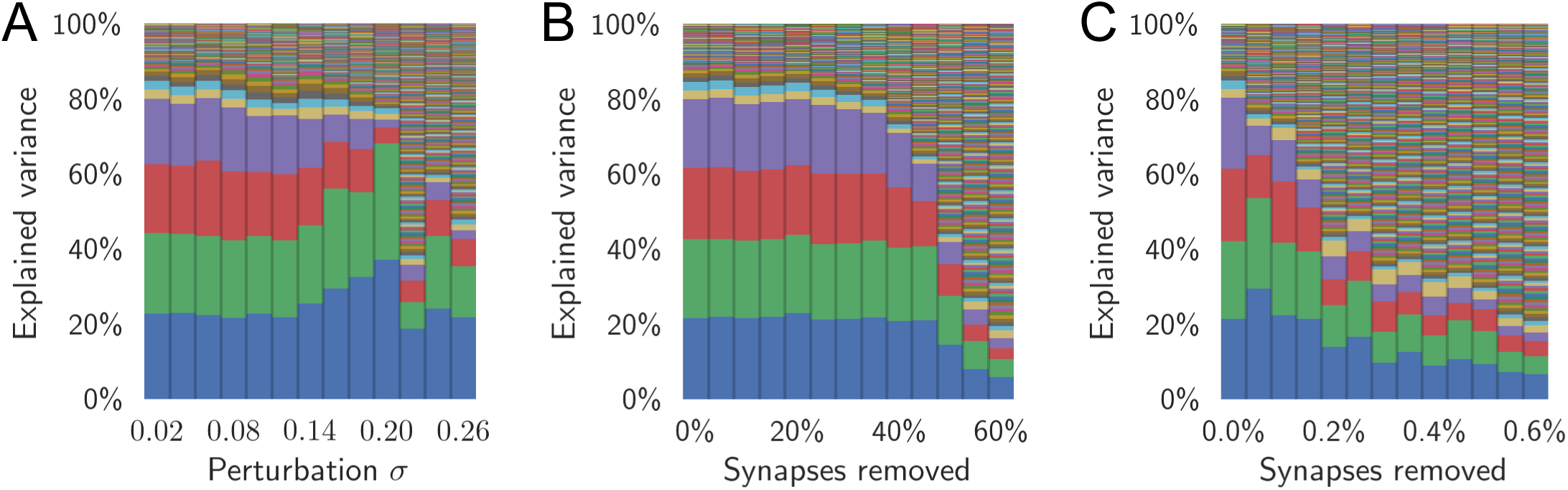
Network resilience to weight perturbations. (**A**) Number of components required to explain the variance (dimensionality) of the population activity changed as the synaptic weights are perturbed. The size of perturbation to the weight matrix was systematically increased. Each synapse was perturbed by an amount *σW_ij_* Each bar corresponds to one such simulation and shows how many components are required to explain the variance (or the dimensionality) for a given size of perturbation. (**B**) Similar to A, but instead of being perturbed synapses are removed in increasing order of absolute weight. The manifold is stable even when almost half of the synapses are removed. (**C**) Similar to B, but instead synapses are removed in decreasing order of absolute weight. Removing the largest synapses results in much greater impairment.

Our simulations showed that as long as the synaptic weight are scaled within 10% of their original values, the distribution of explained variance onto the four dimensions remains stable. As expected, the within-attractor dynamics collapsed before the low dimensionality itself. By collapse of the within-attractor dynamics, we mean that the four Active currents were no longer following the prescribed dynamics. The four fictive currents were designed to be linearly independent so that four dimensions would be clearly visible. Thus, when that design stopped working, the fictive current tend to become partially correlated. Therefore, at about *σ* = 0.20 in Fig. 6A, the dimensionality appears closer to two than four.

## The low-dimensional activity manifold constrains learning

Even though Φ and *K* are not directly visible in the weight matrix, it is useful to consider these variables to understand how the intrinsic manifold of the neuronal activity may constrain learning. While *K* describes the relation between the fictive currents and the synaptic *input*, Φ describes the relation between the filtered *output* and the fictive currents. Due to the somatic and other non-linearities, the optimal Φ does not necessarily equal *K*^T^. However, the transfer functions are typically linear enough for the approximation Φ ≈ *K^T^* not to be unreasonable (Eliasmith and Anderson, 2003, p. 55-56, 191). Indeed, some authors (e.g. Boerlin et al., 2013) have successfully employed this approximation in their network design.

As discussed above, when performing a dimensionality reduction on the spiking activity, the fictive currents are not necessarily perfectly aligned to the projection of the activity to the low dimensional space. We denote this projection Λ ∈ ℝ^*N*×*D*^ for a *D*-dimensional dimensionality reduction. By definition, the columns of Λ consist of the first *D* principal components of the spiking activity. Λ is not strictly equal to *K* nor to Φ*^T^*. However, assuming the neuron transfer functions and binning of spike trains in discrete time bins do not have a prominent effect on the output dimensionality (suggested by Fig. 1), we can argue that the subspace spanned by Λ ought to approximate the subspace spanned by *K*.

In view of these observations we turn our attention to the recent study by Sadtler et al. (2014), and address the question why animals find it hard to learn BCI mappings which lie outside the intrinsic manifold of the network activity? In other words, why the intrinsic structure of the network activity pose strict limits over learning. Sadtler et al. (2014) trained monkeys to control a cursor on a computer screen using the spiking activity recorded from primary motor cortex. In particular, the velocity of the cursor was determined by projecting a sample of the cortical activity down to a two dimensional velocity vector. By changing the projection of the neurons in the mapping, the authors forced the monkeys to relearn the projection, or more precisely, to generate a new network activity that could move the cursor in the desired direction. They found that when the change in the activity projections was designed such that the desired neuronal activity remained in the *intrinsic manifold* — the subspace spanned by Λ in our terminology — the monkeys were able to quickly learn the new mapping and control the cursor. On the other hand, when the activity projection for the BCI task was designed such that the desired activity was outside the intrinsic manifold, the monkeys found it hard to learn the new mapping and in some cases failed to learn.

Using the approximation Λ ≈ *K* ≈ Φ^*T*^, we propose an explanation for the differences in learning performance between inside- and outside-manifold perturbations. An inside-manifold perturbation, as described by Sadtler et al. (2014), can be written as some rotation (possibly including reflection) of the columns of Λ:

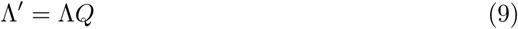

where *Q* ∈ ℝ^*D*×*D*^ is an orthogonal matrix and Λ′ ∈ ℕ × 𝔻 is the new directions of the axes of the subspace. To create this new subspace, assuming Λ ≈ *K*, the monkeys have to change the projection from the fictive currents to the synaptic currents to:

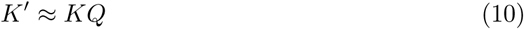

and similarly for the projection from filtered spike trains to the fictive currents:

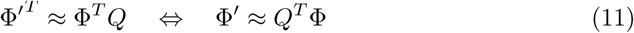

The new weight matrix can then be written as

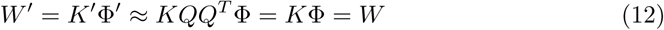

where *QQ^T^* = *I* follows from the fact that *Q* is orthogonal. That is, Eq.12 suggests that inside manifold projections require that *W′* ≈ *W*. This suggests that a relatively small amount of learning may be sufficient to rearrange the axes of the intrinsic manifold.

By contrast, an outside-manifold perturbation requires:

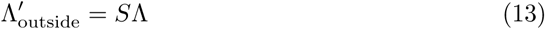

for some orthogonal matrix *S* ∈ ℝ^*N* × *N*^. Following the same steps as above, the weight matrix W′_outside_ required to learn the outside-manifold projections is given by:

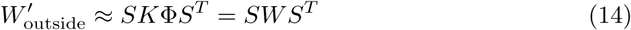

In the generic case, *W′*_outside_ is not equal to *W*. That is, learning *W′*_outside_ to elicit Λ′_outside_ may require substantial synaptic changes and extensive learning.

To support our theory we performed inside- and outside-manifold perturbations on our network model. Specifically, we took inspiration from the procedure Sadtler et al. (2014) used to choose the perturbations and let *Q* be a permutation matrix. However, while Sadtler et al. (2014) used *D* = 10, we kept our choice as *D* = 4. There are 4! = 24 permutations matrices with dimensions 4 × 4, of which one is the identity matrix. Each of these corresponds to a permutation of the columns of *K*. For outside-manifold perturbations, we similarly let *S* be a permutation matrix. This transformation permuted the rows of *K*. However, to limit the the number of possibilities of *S*, we partitioned the rows of *K* into *D* = 4 blocks, as shown in Fig. 7A. We then restricted *S* to permutations of the blocks, rather than of each row individually, resulting in only 4! = 24 possible permutations. Partitioning the blocks this way is similar to how Sadtler et al. (2014) performed the outside-manifold perturbations.

**Fig 7.**
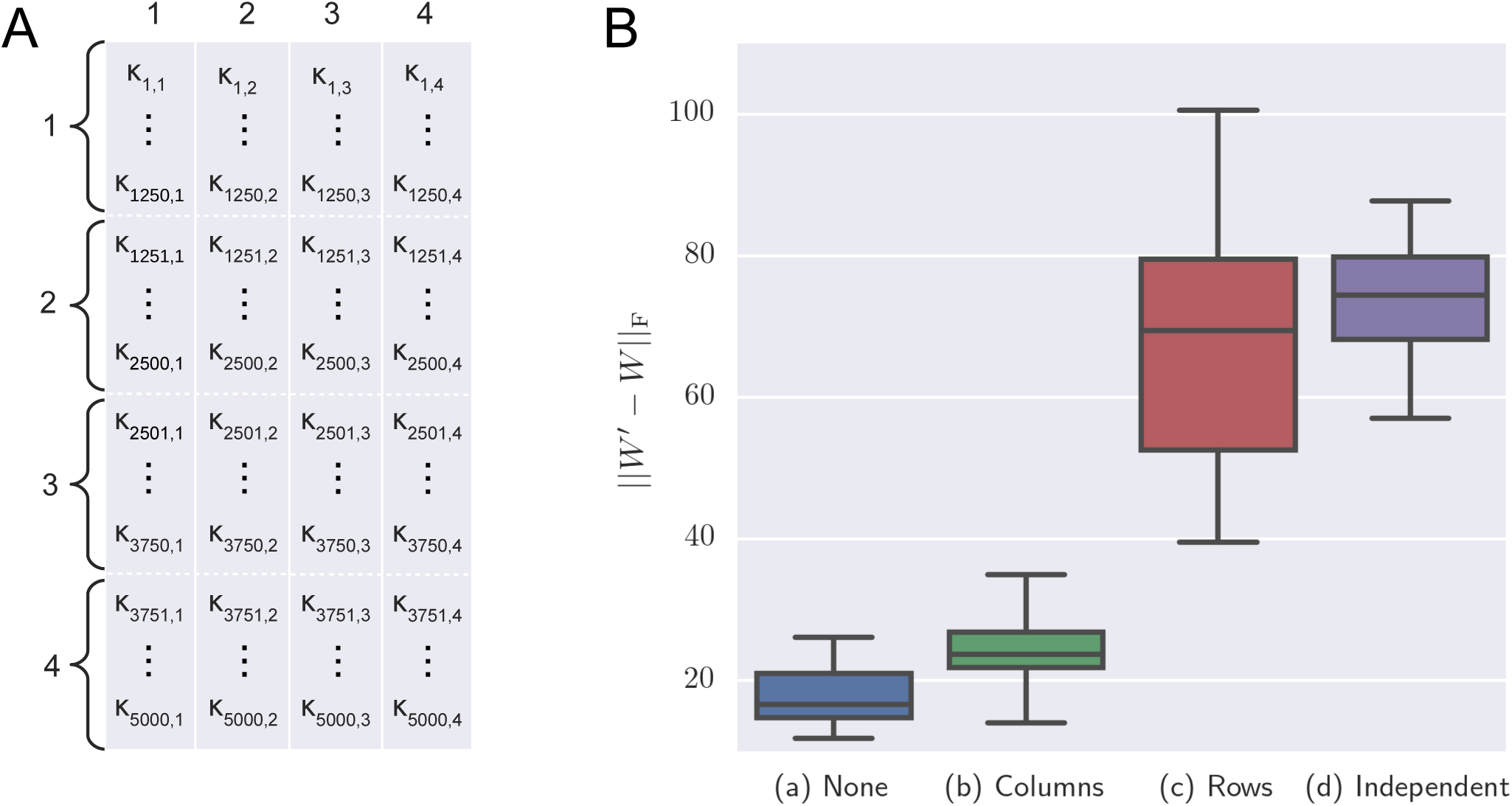
Inside- and outside-manifold permutations. (**A**) In our experiments the dimensions of *K* was 5000 × 4. (**B**) We changed the elements of *K* following four different rules — (**a**) no change at all, (**b**) permuting the four columns, (**c**) grouping the rows into four blocks, and permuting the blocks and (**d**) choosing an independent *K′*. We applied each rule 23 times, going through every permutation for **b** and **c**. For each modification of *K*, we calculated the corresponding weight matrix *W′* and measured its Frobenius difference from *W*. The large Frobenius distance indicates that more weights have to be modified to transform the matrix *W* to *W′*.

For all 23 non-identity permutations, we simulated the network twice: in the first instance we permuted the columns of *K* and in the second instance we permuted the blocks of rows of the matrix *K*. Given the altered *K′* in each simulation we calculated a new *W′* using the linear least squares optimization described above. In other words, we did explicitly include the approximation from Eq. 10, but not from Eq. 11. For comparison, we also simulated the network 23 times without any permutations of *K*, and 23 times where we chose a completely independent *K′*. Because some randomness was introduced in the choice of evaluation points and the initialization of the solver, the weight matrix was not necessarily identical even when *K′* = *K*. For each of these simulations, we compared *W′* to the original *W* (Fig. 7B). As expected from Eq. 12, a permutation of the columns on average resulted only in a marginally different *W′* (Fig. 7Bb). In fact, the change was so small that it was similar to the case when we did not permute the columns (Fig. 7Ba). Permuting the rows (Fig. 7Bc), on other hand, resulted in almost as large change of *W* as if *K′* was chosen randomly (Fig. 7Bd). Thus, these results are consistent with our theoretical explanation that changing the intrinsic manifold of neuronal activity requires large changes in the weight matrix and, therefore, animals may find it hard to learn tasks that involve such transformation of neuronal activity.

The above description shows that the network connectivity (*W*) defines the intrinsic manifold and the dynamical structure of neuronal activity. There is only a small distance between the original network connectivity *W* and a new network connectivity *W′* that remaps the neuronal activity within the original manifold (Fig. 7Bb). That is, only minor adjustments in the synaptic weights may be sufficient to remap the neuronal activity within the intrinsic manifold. This could explain why animals quickly learn the BCI mapping that lied within the intrinsic manifold. On the other hand, a network connectivity that remaps the network activity outside the intrinsic manifold is vastly different from the original matrix *W* that defines the intrinsic manifold (Fig. 7Bc). That is, animals need to retune all the synapses within the network to learn a BCI task that requires them to remap the neuronal activity outside the intrinsic manifold. Thus, our model suggests that animals were slow in acquiring BCI tasks that required them to remap the neuronal activity out of the intrinsic manifold, because such changes would entail bringing in large changes in the strengths of all the synapses in the network.

## Discussion

Recent experiments have found that (1) neuronal activity is constrained to a low-dimensional manifold (Cunningham and Yu, 2014) and (2) animals find it difficult to generate activity that lies outside the intrinsic manifold, thereby containing their learning and behavioral repertoire (Sadtler et al., 2014). Thus far, neuronal mechanisms and functional meaning of the low-dimensionality of the intrinsic activity have remained obscure. At its simplest a network with clustered connectivity can generate low-dimensionality and in that model number of clusters would determine the dimensionality of the network activity (Mazzucato et al., 2016; Williamson et al., 2016). However, clustered connectivity is not consistent with the experimental data. Moreover, such a simple model neither provides a functional reason for the low-dimensionality nor explains why it is difficult to move the neuronal activity out of the intrinsic manifold. Here, we addressed these two questions and propose that: (1) Low-dimensional activity structure reflects the problem the network is designed or has evolved to solve i.e. dimensionality of the network activity is directly related to the complexity of the problem a network is solving. (2) The intrinsic manifold arises due to a specific structure of neuronal connectivity and (3) generation of an activity that lies outside the intrinsic manifold requires a big change in the neuronal connectivity. Because such a large rewiring of connectivity either may not be possible or may take a long time, animals will find it difficult to generate activity outside the intrinsic manifold.

Furthermore, we have shown that neuron transfer-function may not affect the dimensionality of the network activity. In addition, we have also provided a systematic way to generate spiking neuronal networks that follow a certain dynamics and with activity confined to a low-dimensional manifold. This method is based on the *Neural Engineering Framework* (NEF Eliasmith and Anderson, 2003). Specifically, we have demonstrated that the NEF can be used create networks with an arbitrary distribution of variance along the first few principal components. This is a more general feature than creating networks where the variance arbitrarily decreases along the principal components, as is typically seen in most naturally occurring low-dimensional systems. To emphasize this general control, we created a network with activity that varied approximately equally in four directions in the state space, but substantially less in any other direction. However, we do not suggest that having the variance equally distributed along four dimensions accurately reflects biological networks. Rather, we argue that networks created by the NEF and similar frameworks can be designed to have *any* distribution of variance along the *D* first dimensions, given that the dynamical system (Eq. 1) is chosen appropriately. In spite of this ability, they have been neglected as a tool for modeling and explaining low dimensionality in neural networks.

While NEF is a powerful method to construct spiking neuronal networks that conform to a specific dynamical system, the resulting networks often produce highly regular spiking activity patterns. We introduced noisy inputs to the neurons to generate irregular spike patterns without affecting the dynamics of the network activity much. Our approach does not strictly depend on the NEF, and other similar models (e.g. Boerlin et al., 2013; Abbott et al., 2016) could also be used to construct spiking neuronal network that exhibit low-dimensional activity. Unlike NEF, one common feature of the other models is that there are an additional set of connections between neurons in the population that is not subject to the same explicit construction rule (Abbott et al., 2016). In Fig. 2, this would correspond to having lateral connections within the population independently of Φ and *K*. These connections can be either random (e.g. Hoerzer et al., 2012; Sussillo and Abbott, 2009) — essentially turning the network into a *reservoir* or echo state — or specifically designed to optimize the spike times for ease of decoding (e.g. Boerlin et al., 2013). When properly tuned, such extra weights can increase the irregularity and hence the realism of the spike trains. Intuitively, it follows that the induced distortion might also jeopardize the low dimensionality. Surprisingly however, the activity of such networks is in fact low dimensional (Hoerzer et al., 2012; DePasquale et al., 2016). Thus, the matrix Φ could be generated by such approaches without affecting our results, but it remains to be shown and could be a focus of future work.

Our approach to generate spiking neuronal network with low-dimensional activity involves estimation of network weight matrices using a fitting procedure. This, however does not mean that synaptic weights must have an arbitrary precision to generate the prescribed dynamics for a particular network or that there is only one such weight matrix. In fact, the resulting dynamical system is robust to synaptic noise and each synapses can be perturbed by up 20% of its weight before the network loses its intrinsic dimensionality (6A). Moreover, several properties of the the resulting weight matrix are consistent with experimental measurements of the network connectivity e.g. heavy-tailed distribution of synaptic weight, silent synapses and balance of excitation and inhibition for each neuron. Still, the question arises whether such weight matrices and/or dynamical systems can be *learned*? This is a known critique of the NEF and similar frameworks, to which some tentative solutions have been suggested (MacNeil and Eliasmith, 2011; Eliasmith, 2013).

One of the key implications of this model is that knowing the weight matrix W alone is not sufficient to infer the dynamical system implemented by the network. While the weight matrix could be factored into Φ̂ and *K̂*, those may not relate to the original Φ and *K* or may not even have the same dimensions. Similar concerns regarding the weight matrix have been earlier expressed (Buonomano, 2009) and it is not a feature specific to our approach or the NEF. Although knowing the rank of the weight matrix may give hints about the dimensionality of the network activity, we have shown that the weight matrix can have considerably higher rank than the activity itself. However, that information can also be inferred from the spiking activity itself which is much simpler to measure. In fact, the low dimensionality implies that sampling of only a few neurons may be sufficient to get a good estimate of the dimensionality. In addition, if our model is correct, the dimensionality of the network activity is also visible in the input to the neuron and can also be inferred from the sub-threshold membrane potential or calcium imaging signals.

Next, when the weight matrix encodes a dynamical system, the dimensionality of the resulting network activity is independent of the number of neurons sampled from the network as long as the count of sampled neurons is larger than the dimensionality of the underlying dynamics system. Unlike previous suggestion, clustered connectivity is not necessary to keep the dimensionality low and independent of the sample size. In our model the synaptic weights exhibit a heavy tailed-distribution (Fig. 3C). The weak synapses, however do not contribute much to the resulting dynamics and up to 40% of the weakest synapses can be removed without affecting the dynamics (Fig. 6B). By contrast, the network dimensionality and dynamics are more susceptible to removal of the strong synapses (Fig. 6C). Thus, the model predicts that a loss of a few strong synapses may be detrimental to the network function.

Throughout this text we have emphasized on designing network with low-dimensional dynamics. Here, and in other similar literature low-dimensionality refers to the fact the number of components required to explain the variance of the network activity are far smaller than the number of neurons in the network. That is, whether a dimensionality *D* is small or not depends on the number of neurons in the network. Here, we have extended that concept one step further and have suggested that the dimensionality of a network activity refers to the number of variables a network control while performing a certain task, for instance to implement a dynamics system. It can be argued that because tasks such as sensorimotor transformation and working memory are inherently low-dimensional, the resulting activity in these tasks is also low-dimensional. More complex cognitive tasks may exhibit higher dimensionality Fusi et al. (2016). In addition, it is also possible that the low-dimensional dynamics is only a feature of a specific neuron type and inclusion of other cell-types may increase the dimensionality.

## Acknowledgements

We would like to thank Ylva Jansson and Jan Pieczkowski for helpful discussions and comments on the manuscript. Partial funding from the School of Computer Science and Communications, KTH, Parkinsonfonden Sweden and The Strategic Research Area Neuroscience (StratNeuro) program coordinated by Karolinska Institutet Stockholm, Sweden is greatly acknowledged.

